# Short-Delay Neurofeedback Facilitates Training of the Parietal Alpha Rhythm

**DOI:** 10.1101/2020.07.27.222786

**Authors:** Anastasiia Belinskaia, Nikolai Smetanin, Mikhail Lebedev, Alexei Ossadtchi

**Affiliations:** Center for Bioelectric Interfaces, Higher School of Economics, Moscow, Russia, 101000

## Abstract

The therapeutic effects of neurofeedback (NFB) remain controversial. Here we show that visual NFB of parietal electroencephalographic (EEG) alpha-activity is efficient only when delivered to human subjects at short latency, which guarantees that NFB arrives when an alpha spindle is still ongoing. NFB was displayed either as soon as EEG envelope was processed, or with an extra 250 or 500-ms delay. The time course of NFB-induced changes in the alpha rhythm clearly depended on NFB latency, as shown with the adaptive Neyman test. NFB had a strong effect on the alpha-spindle incidence rate, but not on their duration or amplitude. The sustained changes in alpha activity measured after the completion of NFB training were negatively correlated to latency, with the maximum change for the shortest tested latency and no change for the longest. Such a considerable effect of NFB latency on the alpha-activity temporal structure could explain some of the previous inconsistent results, where latency was neither controlled nor documented. Clinical practitioners and manufacturers of NFB equipment should add latency to their specifications while enabling latency monitoring and supporting short-latency operations.

## Introduction

In this study, we implemented a neurofeedback (NFB) task, where participants self-controlled their parietal electroencephalographic (EEG) alpha rhythm, and investigated how setting NFB latency to different values affected EEG patterns.

NFB is a closed-loop paradigm, where subjects are presented with an indicator of their own brain activity, which they learn to change in a certain desired way [25], [26], [53], [55], [60]. In a typical NFB experiment, neural activity is recorded, converted into features of interest, processed to generate an output signal, and delivered to the subject as a sensory stimulus, typically visual or auditory. Training with NFB results in plastic changes in the neural circuits involved. This type of learning fits the definition of operant conditioning, where the behavioral responses are derived directly from neural activity, and indicators of successful performance presenting to the subjects serve as rewards that reinforce the wanted behavior [53]. Historically, the first implementations of NFB were based on EEG recordings [25], [26], followed by more recent demonstrations based on such recording methods as magnetoencephalography (MEG) [38], [4], functional magnetic resonance imaging (fMRI) [59], [65], functional near-infrared spectroscopy (fNIRS) [28], and depth electrodes [68].

Practical interest to NFB approach is driven by the expectation that it could become a powerful therapy that improves brain processing in subjects suffering from neurological disorders [60]. Since, at least theoretically, NFB could target neural circuits very specifically, this method could supplement or even replace the traditional pharmacological [63], [3], [70], [31] and cognitive-enhancement [69], [10] therapies. However, despite the high expectations and numerous studies on NFB-based therapies, this approach remains controversial because of the high variability of its outcomes and the lack of improvement in a considerable number of cases [2], [60]. The difficulties in the development of efficient NFB-based therapies are rooted in the insufficient understanding of the physiological effects of NFB [24], issues related to ergonomics, and problems in signal processing [22].

Here we looked into the issue that has not been sufficiently addressed by previous research on NFB: the proper setting of NFB latency, that is the time interval from the occurrence of a neural activity till the delivery of the feedback of that activity to the subject. NFB latency specifies the reinforcement schedule [53] and as such it should significantly affect the outcome of operant conditioning [49], [44].

Temporal specificity is of essential importance for neural processing [23], and feedback latency plays a pivotal role in a range of closed-loop systems, including nonlinear dynamical systems [66] and biological systems with a delay [33], [6]. Moreover, human psychophysics studies have demonstrated that feedback latency significantly influences sensory, motor, and cognitive processing. Thus, visual perception and performance are impaired when human subjects observe images on the displays with lags and slow frame rate [9]. Additionally, delaying visual feedback diminishes the accuracy of drawing [17], and accuracy of slow but not fast reaching movements toward a target is impaired if the light is suddenly turned off [27]. In a virtual environment, feedback latency affects the sense of presence, particularly when the environment is stressful [35]. Furthermore, the sense of agency, that is the perception of being in control of own movements, deteriorates with an increase of the visual feedback delay [16]. Similarly, the telepresence level, when using a surgical robot, decreases with increasing feedback delay [50].

The effects of feedback latency have been recognized in the literature on NFB and brain-computer interfaces (BCIs). Oblak et al. [37] simulated fMRI signals in visual cortex and had human subjects develop cognitive strategies to utilize NFB produced from the simulated data. They observed a better performance for continuous NFB than for intermittent NFB. Additionally, NFB delay significantly affected the performance with continuous NFB. Furthermore, they developed a computational model of automatic NFB learning. This modeling showed that when NFB was blurred and arrived with a delay, the performance was better with intermittent feedback compared to continuous feedback. The authors suggested that NFB settings should be optimized to match the experimental paradigm and learning mechanism (cognitive versus automatic). Evans et al. [14] implemented a motor imagery-based BCI where the control signal was derived from sensorimotor mu or beta rhythms. They showed that introduction of a feedback delay resulted in a reduced sense of agency, that is subjects did not perceive the BCI output as the result of their voluntary intentions.

In the present study, we implemented a NFB derived from the parietal alpha rhythm and for the first time systematically investigated the effect of NFB latency on the changes in EEG patterns. There is a vast literature on brain rhythms [7] and on NFB paradigms for controlling the rhythms in different brain areas [69], [18], [58]. Brain rhythms typically wax and wane, which makes it important to understand their temporal structure when setting NFB parameters. In addition to the temporal dimension, space [32] and frequency [20], [62] are the dimensions that carry specific information that could be utilized to generate NFB. The following reasons motivated our choice of alpha rhythm as the source of NFB. First, alpha rhythm is one of the most prominent and most responsive to training brain rhythms [36], [1], [12], [19], [21], [61]. According to the existing literature and our own results [41], parietal alpha rhythm is easy to isolate (in contrast to the sensorimotor rhythm, for example) and easy to train with NFB practically in all subjects. Moreover, NFB that is based on the parietal alpha rhythms has been suggested as an approach to gaining a range of functional improvements, including improvements in cognition [61], [1], [19], attention [4], [39], [40], [8], working memory [11], [67], mood [42], [13], [43], and relaxation [5].

NFB latency results from the signal processing pipelines that comprises several stages, see Figure 1 A. The overall latency is the time from the occurrence of a neuronal event being tracked till the instance when the subject receives an NFB replica of this event. This time includes a data collection interval (tens of milliseconds), interval of filtering and instantaneous power estimation (hundreds of milliseconds), and feedback generation interval (tens of milliseconds). In the currently used NFB systems, latency falls within the range from 300 to 1000 ms. This duration can be decreased by means of optimizing data acquisition and signal processing steps. To this end, we recently developed NFB-Lab software [57] that supports a novel causal complex-valued finite impulse response (cFIR) approach for simultaneous narrow-band filtering and extracting the instantaneous power of an EEG rhythm [56]. This method allowed us to achieve high accuracy of narrow-band power estimation with the lag as short as 100 ms. In this implementation, communication with the EEG recording system was enabled by the Lab Streaming Layer protocol [29], followed by prefiltering, and visual stimulus delivery which all together incurred approximately a 144-ms average lag. Thus, the overall latency was 244 ms in our system. We refer to this delay as system’s base latency.

**Figure 1:**
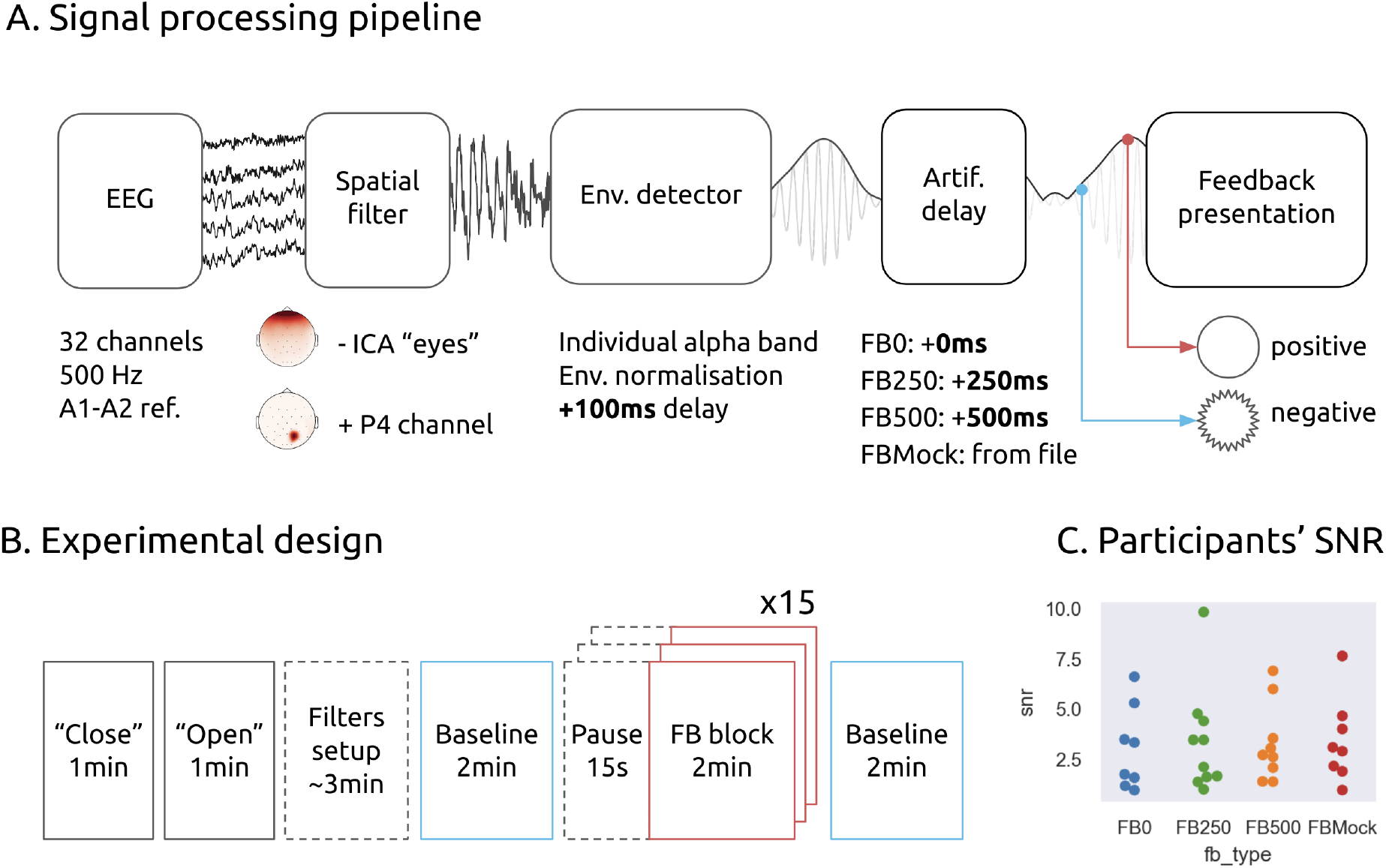
Schematics of the experiment and signal processing. A: Signal processing pipeline, where multichannel EEG signals are processed by a spatial filter to remove artifacts, converted into a narrow-band signal, time shifted with an artificial delay, and converted into a visual feedback. B: Experimental design, where resting-state EEG activity is recorded first, followed by NFB sessions. At the end, the post-training baseline is recorded. C: Signal to noise ratio of the alpha rhythm across NFB groups. Datapoints represent SNR value for each subject.

In this study, we used the NFB Lab software and the cFIR approach to examine NFB mechanisms for several overall latency values in the 244–744 ms range. We found that delayed NFB impeded both transient changes in EEG patterns that occurred during training and the changes sustained after the training was completed.

## Methods

### Participants and settings

Forty healthy right-handed subjects (13 males and 27 females; aged 24.58 +- 5.3 years, mean +- SD) participated in the experiments. The study was conducted in accordance with the ethical standards of the 1964 Declaration of Helsinki. All participants provided written informed consent prior to the experiments. The ethics research committee of the National Research University, The Higher School of Economics approved the experimental protocol of this study.

The NFB signal was derived from the P4 channel (corresponding to the right parietal region). An alpha envelope was visualized as a circle with a pulsating outline. The subjects had to smooth that outline by increasing their P4 alpha-band power.

All participants were instructed to refrain from using any conscious strategy. This assured that learning to control the parietal alpha rhythm was automatic [30], that is the mode where the latency of a continuous NFB has the strongest effects [37].

Subjects sat in a comfortable chair at a distance of 80 cm from an LCD monitor with a 24-cm diagonal and a 60-Hz refresh rate. These settings remained constant throughout the entire experiment.

EEG signals were recorded using 32 AgCl electrodes positioned according to the 10-20-system. Each EEG channel was sampled at 500 Hz using an NVX-136 amplifier (Medical Computer Systems Ltd), and bandpass-filtered in the 0.5 - 70 Hz band. These preprocessing filters incurred an overall delay of no more than 10 ms for the EEG bandwidth of interest (8–12 Hz). Digital common ear reference was derived from the electrodes placed on both ears. The impedance for each electrode was kept below 10 KOhm.

### Experimental protocol and measurements

The participants were split into four equal groups, each with different NFB settings: (1) NFB with no delay added to the system’s base latency of 244 ms (FB0), (2) NFB with a 250-ms delay added (FB250), (3) NFB with a 500-ms delay added (FB500), (4) mock NFB where the feedback was generated from EEG data taken from a different participant (FBmock). Due to a technical problem associated with an intermittently suspended EEG-software communication, we could not use the records of 5 subjects (2 from FB0 group, 2 from FBMock group, and 1 from FB500 group); these data were excluded. When choosing the statistical methods, we took into consideration these unequal sample sizes. The details of the statistical analysis can be found in the section Results and Appendix: Comparing learning curves.

The experimental sequence is shown in Figure 1 B. Prior to NFB sessions, we recorded resting-state EEG and used these data to set the spatial filters for the suppression of eye-movement artifacts and determine the frequency of alpha rhythm in individual subjects (for more details, see EEG data processing). Next, just prior to NFB training, we recorded a 2-min baseline with eyes open. Then, NFB training started that comprised fifteen two-minute blocks of NFB separated by 15-s resting periods. Immediately following the NFB training, we recorded the 2-minute post-training baseline.

### Composition of subject groups

When composing the subject groups, we had to deal with the differences in alpha-rhythm patterns in individual subjects. These individual features were evident in the resting-state data. It was previously shown that resting-state alpha amplitude is predictive of a subject’s subsequent improvement in NFB control with training [64]. This effect could contribute to the heterogeneity of subjects across groups (FB0, FB250, FB500, FBmock) in our study. For instance, a subject with a weak alpha rhythm would not be able to improve in NFB control in any condition, but his/her assignment to a particular group could produce a false group-related result. To control for this effect, we implemented a stratified sampling procedure [47], [45], [51], [48] that equalized resting-state alpha amplitude across groups. We measured signal to noise ratio (SNR) for the alpha amplitude while participants rested with open eyes, and categorized them as having high alpha (SNR >4,4; Mean (2,89) +SD (1,51)), low alpha (SNR <1,38; Mean (2,89) - SD(1,51)), or medium alpha (1,38 <SNR <4,4). Each subject was assigned to a NFB group with the procedure where the NFB group was randomly selected, but if the SNR category (high, low or medium alpha) was already filled for that group with the entries from the other subjects, random selection repeated [47] (for more details, see “Signal to noise ratio” section below). Wilcoxon rank-sum test showed no statistical difference between the SNR across the four experimental groups (Figure 1 C).

### EEG data processing

#### Independent component analysis

As shown in Figure 1 B, the experiments started with the recording where a subject first looked at the fixation cross for one minute and then closed the eyes for one more minute. We then used these data to build a spatial filter based on the independent component analysis (ICA) to remove the artifacts caused by eye movements and blinking. This approach decomposed the EEG signals into independent components, including the ones containing the artifacts. Ocular artifact components were detected as those with the largest value of mutual information of their time-series and the signals in Fp1 and Fp2 channels, which are closest to the eyes. A spatial filter matrix was then constructed for the subsequent online application during the NFB sessions.

#### Individualized bandpass filter

Bandpass filters for extracting alpha activity were built separately for each individual. The signal was taken from channel P4. To detect alpha activity, we started with the frequency interval from 8 to 12 Hz, and then adjusted the interval and filter parameters for each individual subject. Namely, we determined the central frequency *F_c_* of the rhythm by visual inspection and then set the interval to [*F_c_* – 2, *F_c_* + 2].

#### Signal to noise ratio

The SNR was calculated before the main session from the two-minute baseline as the ratio of the average power spectral density (PSD) magnitude within the individually determined target frequency range *[F_c_* – 2, *F_c_* + 2] to the mean magnitude of PSD within the two flanker sub-bands: [*F_c_* – 4, *F_c_* – 2) and (*F_c_* + 2, *F_c_* + 4]. The participants with SNR less or equal to 1 were not included into the pool of subjects. The participants with SNR greater than 1 were assigned to one of the experimental groups with a stratified sampling procedure (see “Composition of subject group” section above).

#### Envelope extraction

Alpha-rhythm envelope was extracted with the cFIR approach [56] (Figure 2 A). In this method, the raw EEG signal is transformed into a narrowband analytic signal, a complex-valued function whose absolute value corresponds to an instantaneous amplitude(or envelope) of the rhythm. The cFIR method explicitly defines NFB latency and obtains a more accurate envelope estimate for a specified latency compared to the other approaches to quantification of narrowband components in the EEG data, [56]. This speed-accuracy trade-off can be appreciated from the accuracy vs. processing delay curves presented in Figure 2 B for the cFIR and the commonly used approach based on narrow-band filtering followed by signal rectification. In the present study, we set the cFIR delay parameter to 100 ms (the point marked by a cross in Figure 2 B). This setting corresponds to the correlation coefficient of 0.85 +- 0.1 between the actual and the on-line reconstructed envelopes.

**Figure 2:**
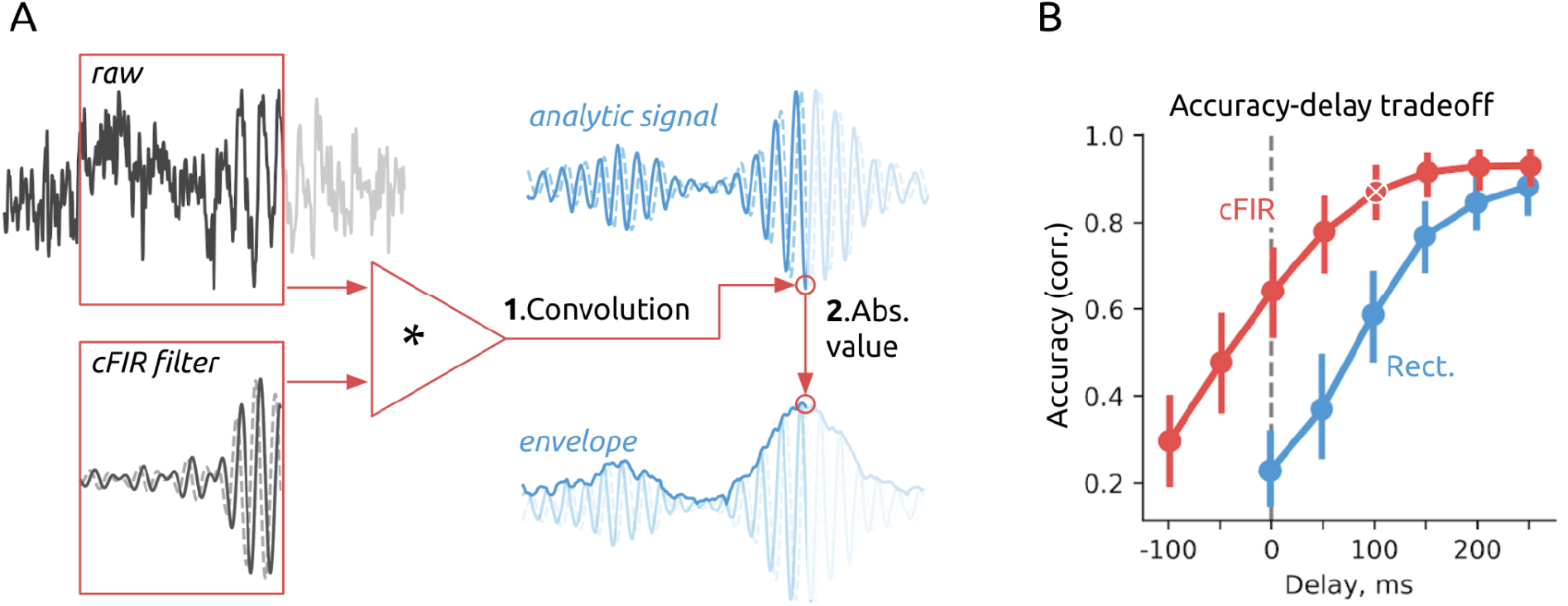
Schematics of envelope extraction. A: Graphical illustration of the cFIR method, where the envelope is obtained from the optimally-filtered complex-valued analytic signal. B: Speed-accuracy curve for envelope extraction with cFIR (red) and Rect (blue) methods. cFIR outperforms Rect for all latency values. The cFIR setting used in this study (latency of 100 ms) is marked by a cross.

#### Latency measurement

To monitor the system’s total latency in accordance with the recommendations of the CRED-nf checklist [46], we used a direct latency measurement aided with a photosensor attached to the corner of the screen [57]. Figure 3 A explains this method for syncing the “brain time” (top) – the actual time of the occurrence of neural events, with the “PC time” – the time when these events are registered on the computer.

**Figure 3:**
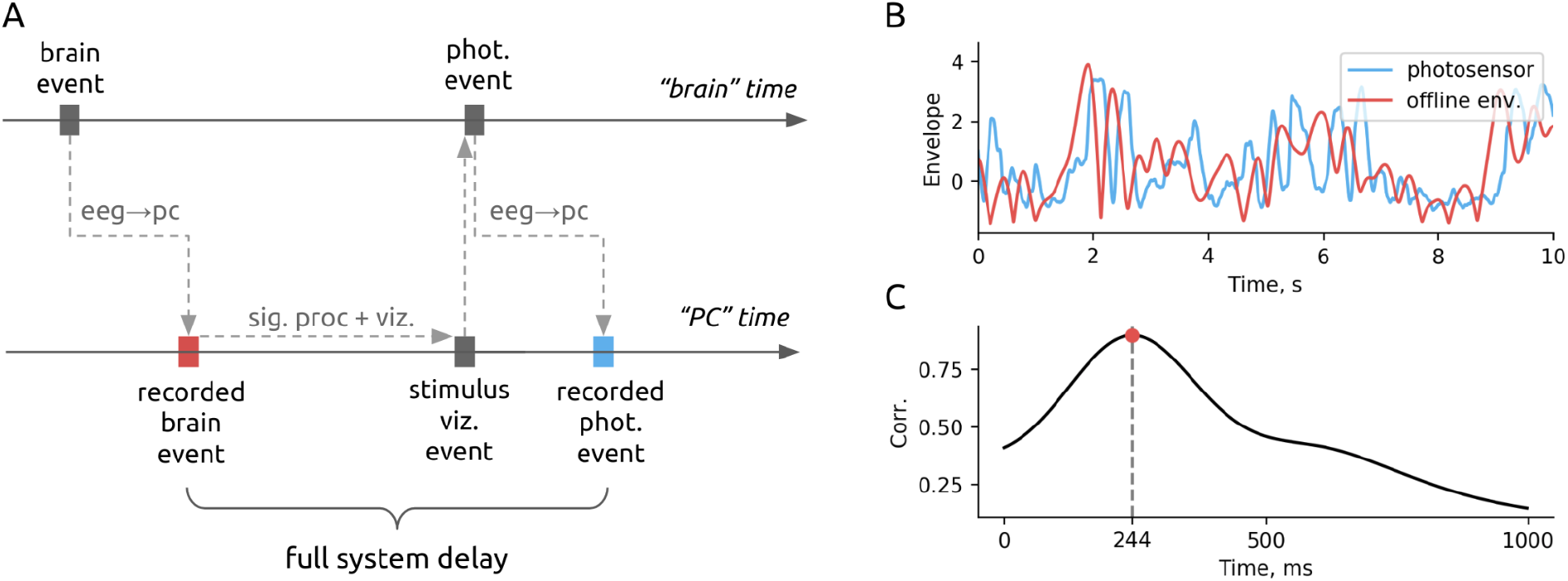
Schematics of the overall latency measurement. A: Syncing of the “brain time” (top) and “PC time” (bottom) using a photosensor. EEG signals (brain events) are registered by the computer with an EEG to PC lag. The computer sends a signal to the screen, which is measured by the photosensor and is then sent back to the computer through an auxiliary EEG channel with the same EEG to PC lag. The overall latency is the time from a brain event of interest (instantaneous alpha-band amplitude) till the corresponding photosensor event, with both measurements referred to the PC time axis. B: Zero-latency alpha amplitude, *y*_0_(*t*) (red) and the real-time NFB captured by the photosensor, *p*(*t*) (blue). C: Crosscorrelation function betweeen *y*_0_(*t*) and *p*(*t*) as a function of the time lag. The time of the crosscorrelation peak corresponds to the overall NFB latency.

EEG signals were sampled by an EEGgraph and sent to the PC, where they were timestamped with an EEG to PC transmission delay. The EEG data were processed and converted into a feedback stimulus to be shown on the screen. The time of the screen event was detected by the photo-sensor whose output was fed to one of the EEGgraph channels and transmitted back to the computer with the same delay as for the EEG to PC transfer. Therefore, the lag between the neural event (in our case, instantaneous alpha amplitude extracted from EEG with zero latency in an offline analysis) and the corresponding photosensor event was measured in reference to the PC time axis: it corresponded to the overall NFB latency.

We implemented this method by dedicating a small square in the upper-right screen corner to the photosensor signal. The square brightness corresponded to the NFB signal presented on the main portion of the screen, and the square itself was covered by the photosensor and therefore invisible to the participants. The square brightness, measured by the photosensor, was fed to an auxiliary channel of the EEG device and recorded together with the EEG data. The overall latency was estimated by comparing the photosensor signal, *p*(*t*), with the zero-latency NFB signal calculated offline, *y*_0_(*t*) (Figure 3B). The lag between these two signals was calculated as the timing of the peak in their cross-correlation function (Figure 3 C). With this approach, amplitude and time, (*R_max_*, *τ_max_*), of the cross-correlation peak were computed for the entire duration of the experiment and used as the measurements of NFB average accuracy and latency.

To test the effects of changing the overall latency, we either used the nominal 244-ms latency (FB0) or artificially added an extra delay of 250 ms (FB250) or 500 ms (FB500).Mock feedback (FBMock) was used as a control condition.

#### Blinding

Theexperiments were designed to have minimal interaction between the experimenter and subject after the subject was assigned to a NFB-latency group. This was assured by running the randomized stratification procedure after setting the individual filters. The group assignment was generated by the computer and saved in the subject’s folder based on the recordings of the first two 1-minute baselines; no interaction between the subject and experimenter occurred at that point.

When processing the data and generating the results, we analyzed the data for all subjects and all groups with a single script that applied the same processing pipeline to all entries. Next, the results of this stereotypical processing were grouped using the group assignment variable and the appropriate statistical comparison was performed automatically. The data and analysis scripts can be found at https://github.com/nikolaims/delayed_nfb

## Results

### NFB latency affects learning curve profiles

We first explored the changes in P4 alpha-band magnitude as a function of training block sequential index. The shapes of these curves are different depending on NFB latency, which points to latency-specific training *dynamics*. Figure 4 shows the across-subject means of alpha-band magnitude computed for 15 training blocks; separate curves correspond to different NFB delays. In order to minimize inter-subject variability, each subject’s data were normalized by dividing by the mean magnitude for all training blocks. With this normalization, a “flat” curve stabilized around the level of 1 would indicate an absence of training effect. If NFB training results in alpha power increase, the across-subject mean values form a curve with an overall positive slope. Figure 4 shows that NFB training resulted in a noticeable gradual enhancement of the alpha rhythm magnitude for all tested latency values, and only a small enhancement occurred for the mock NFB.

**Figure 4:**
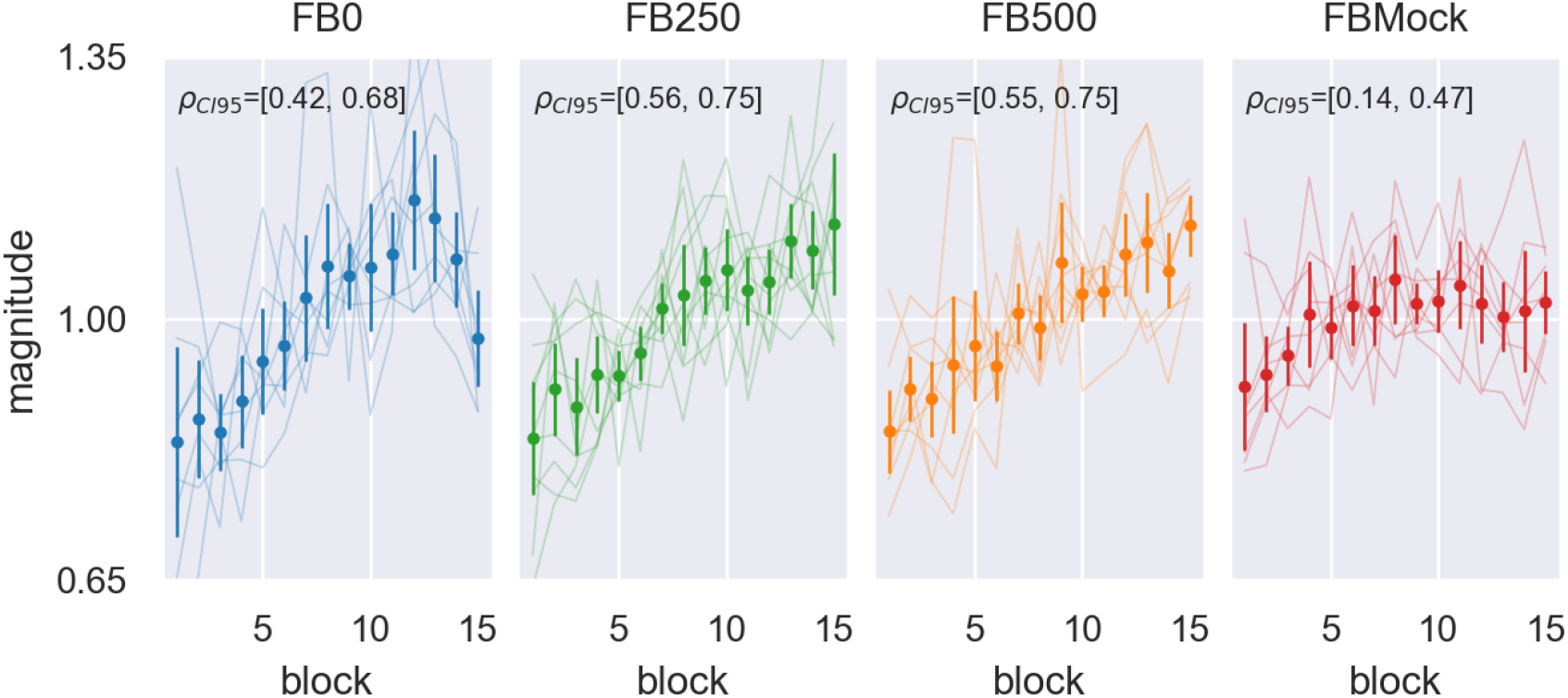
Learning curves reflecting changes in the alpha-band magnitude across 15 experimental blocks. Data for FB0, FB250, FB500, and FBMock conditions are shown in separate panels.Thin lines correspond to individual subjects. Thick lines represent across-subject averages. Vertical bars correspond to two standard errors. The numbers on each plot correspond to the 95 % confidence interval for linear correlation.

We first quantified the changes in alpha magnitude using a classical linear regression model. The confidence intervals (CIs) for the linear correlation coefficient *ρ* are displayed on top of each panel in Figure 4. In the FBMock condition, the learning curve is noticeably flatter compared to the NFB conditions; iowever, in all four groups including FBMock, we found a significantly positive correlation between mean alpha rhythm magnitude within a block and block’s sequential number. The 95 % confidence interval on *ρ* in FBMock condition does not overlap with the CIs for FB250 and FB500 conditions. The CIs for the linear correlation coefficient in FB0 and FBMock conditions overlap very slightly, primarily because of the prominent decline that occurred during the last three blocks in the FB0 condition.

For a more detailed quantification of the learning curves, we performed the adaptive Neyman test (AN-test) proposed in [15]. This test considers the projections of the differences in *t*-statistics onto a set of orthogonal basis functions and assesses the significance of the projection coefficients. This approach takes into account the fact that the observed samples belong to a smooth curve. We applied this test but used Legendre orthogonal polynomials instead of Fourier basis as proposed in the original paper. This was done because the shapes of these individual basis functions allowed for a more parsimonious description of the typical NFB learning dynamics. The details of our implementation of this test can be found in Appendix: Comparing learning curves. Figure 5 A shows the results of applying the AN-test to pairwise comparison of P4 alpha magnitude learning curves observed in the four conditions.

**Figure 5:**
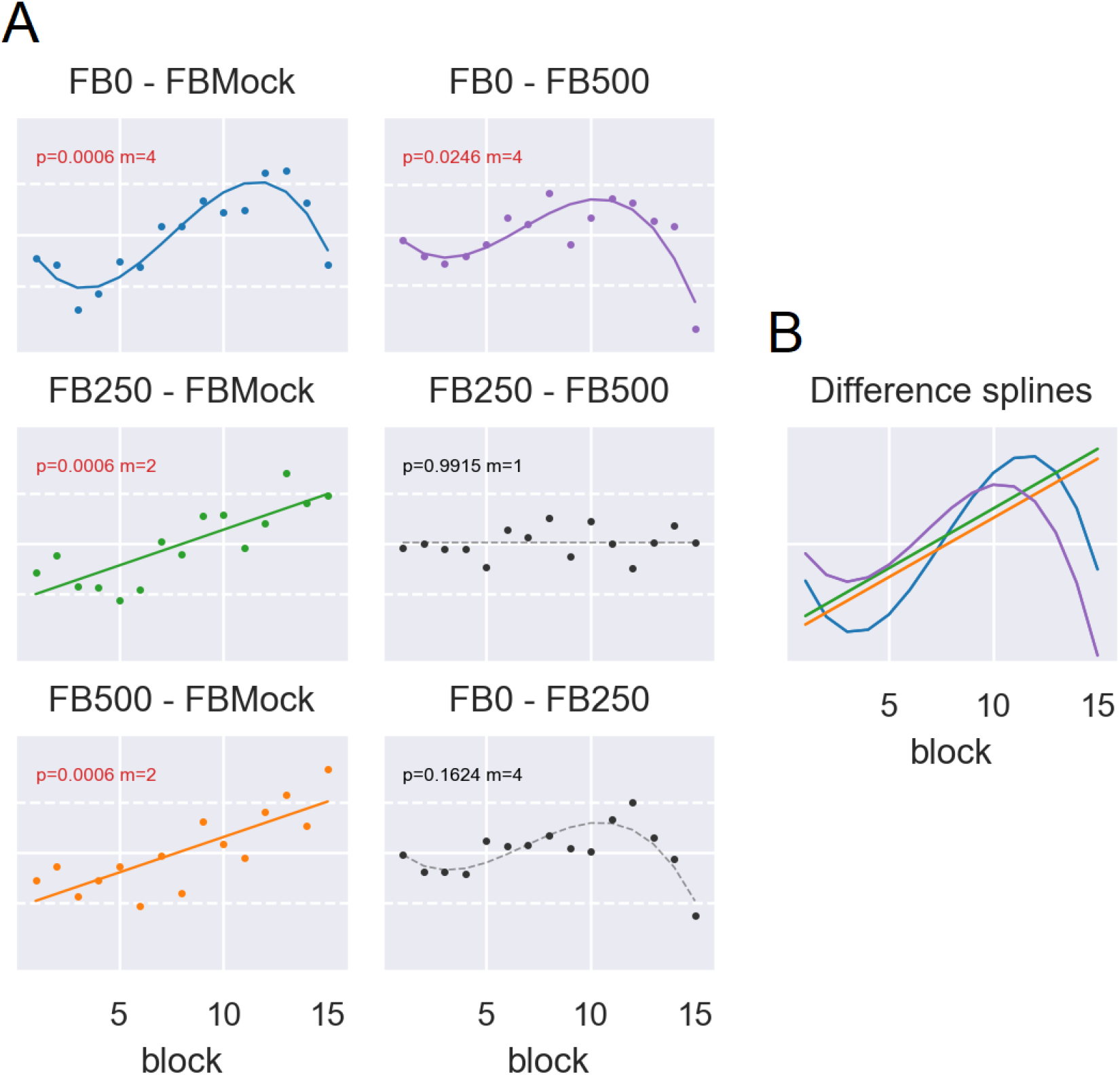
A detailed comparative pairwise analysis of alpha magnitude learning curves observed in four conditions. A: Pairwise *t*-statistics profiles. The corresponding FDR-corrected *p*-values and the number of used coefficients are shown in the top of each plot. B: Parametric differential profiles computed as linear combinations of significant polynomials for the four significant (*p* < 0.05) pairwise comparisons.

As with the simple linear model, we found that all NFB conditions had training dynamics that were significantly different from that observed in FBMock condition. Interestingly, FB0 vs. FBMock *t*-statistics profile is characterized by the S-shape indicating the later onset of positive changes and earlier saturation with the subsequent decline. The test was also powerful enough to detect the difference in the learning curve shapes observed in FB0 and FB500 conditions. As evident from Figure 5 B, FB0 was characterized by a steeper learning curve compared to the ones observed in the other conditions (see FB0–FBMock and FB0–FB500 profiles). While the FB0 curve had the steepest rise, it saturated earlier compared to the other conditions and then declined. EEG rhythmic activity is non-stationary and consists of a succession of transient burst events. Changes in mean magnitude of alpha activity may be caused by variations in burst duration, burst amplitude, and the incidence rate of such bursts.

In our previous study with a relatively short and fixed NFB latency of 360 ms [41], we observed NFB-evoked changes in the incidence rate of alpha spindles but not in their amplitude and duration. Accordingly, alpha spindles can be considered as discrete structural units whose characteristics could change as the result of NFB training. A similar view was expressed in [54], where the functional role was highlighted of discrete beta-spindles for motor control in several species.

Following the methodology of our previous study [41], for all four conditions, we analyzed the time course of spindle characteristics, including incidence rate, amplitude, and duration. Alpha spindles were extracted with simple thresholding from the envelope of the alpha-band filtered P4 time course, see Figure 6. By the analysis design, the characteristics of burst depended on the selected threshold. For example, if a high threshold value was used, only the most prominent episodes of alpha activity were qualified as bursts (Figure 6 A), while with a lower threshold value, the same bursts appeared to have of a longer duration. Additionally, less prominent alpha episodes were also counted (Figure 6 B). Furthermore, with a low threshold, adjacent bursts could merge into a single one, thereby increasing the duration of bursts and reducing their count.

**Figure 6:**
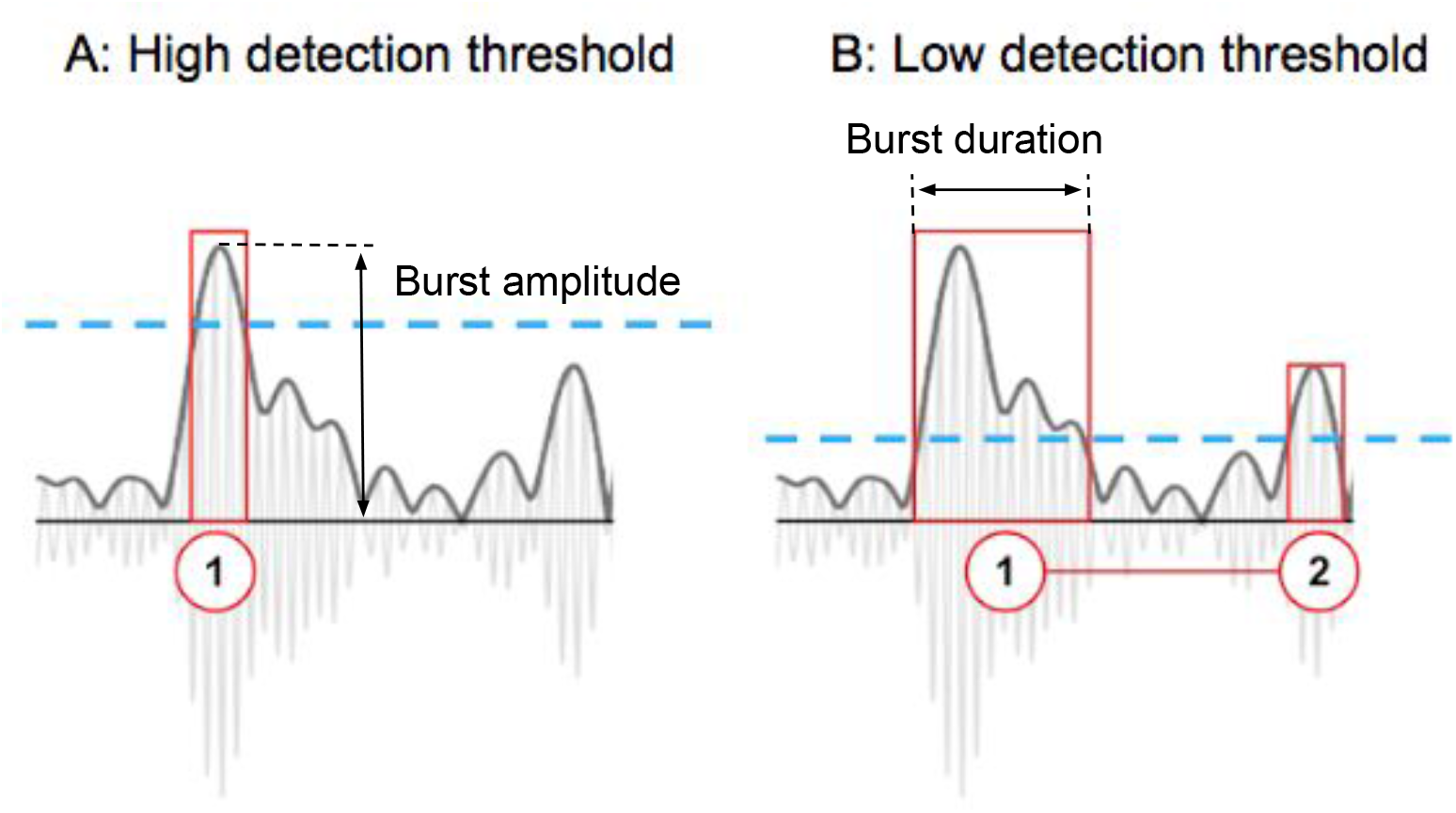
Detection of alpha bursts with different threshold values. Burst amplitude and duration characteristics. The curve represents alpha envelope. A: A single, narrow burst is selected with a high threshold B: The selected burst widens and an additional burst is detected with a lower threshold.

As a reasonable solution, we selected the threshold value that corresponded to the minimum mean mutual information (MI) between the three distinct pairs of morphological parameters: incidence rate vs. burst amplitude, incidence rate vs. burst duration, and burst amplitude vs. burst duration. This analysis leads us to select the threshold factor of *μ* = 2.5. The actual threshold was then found by multiplying this factor by the median value of the time series.

To assess the effect of NFB training on the parameters of alpha bursts, we compared three NFB groups (FB0, FB250, FB500) to FBMock group. The results of the AN-test for pairwise comparison for all rhythm characteristics changes (magnitude, number of bursts, amplitude, and duration) are shown in Figure 7 A. For all NFB conditions, the curves representing burst incidence rate are different from the corresponding curve for FBMock condition (*p* < 0.02, FDR corrected). As to the curves for alpha-spindle duration and amplitude no significant difference is present between the NFB and mock NFB conditions. This result replicates our previous observation [41] and extends it to a broader range of NFB latency values.

**Figure 7:**
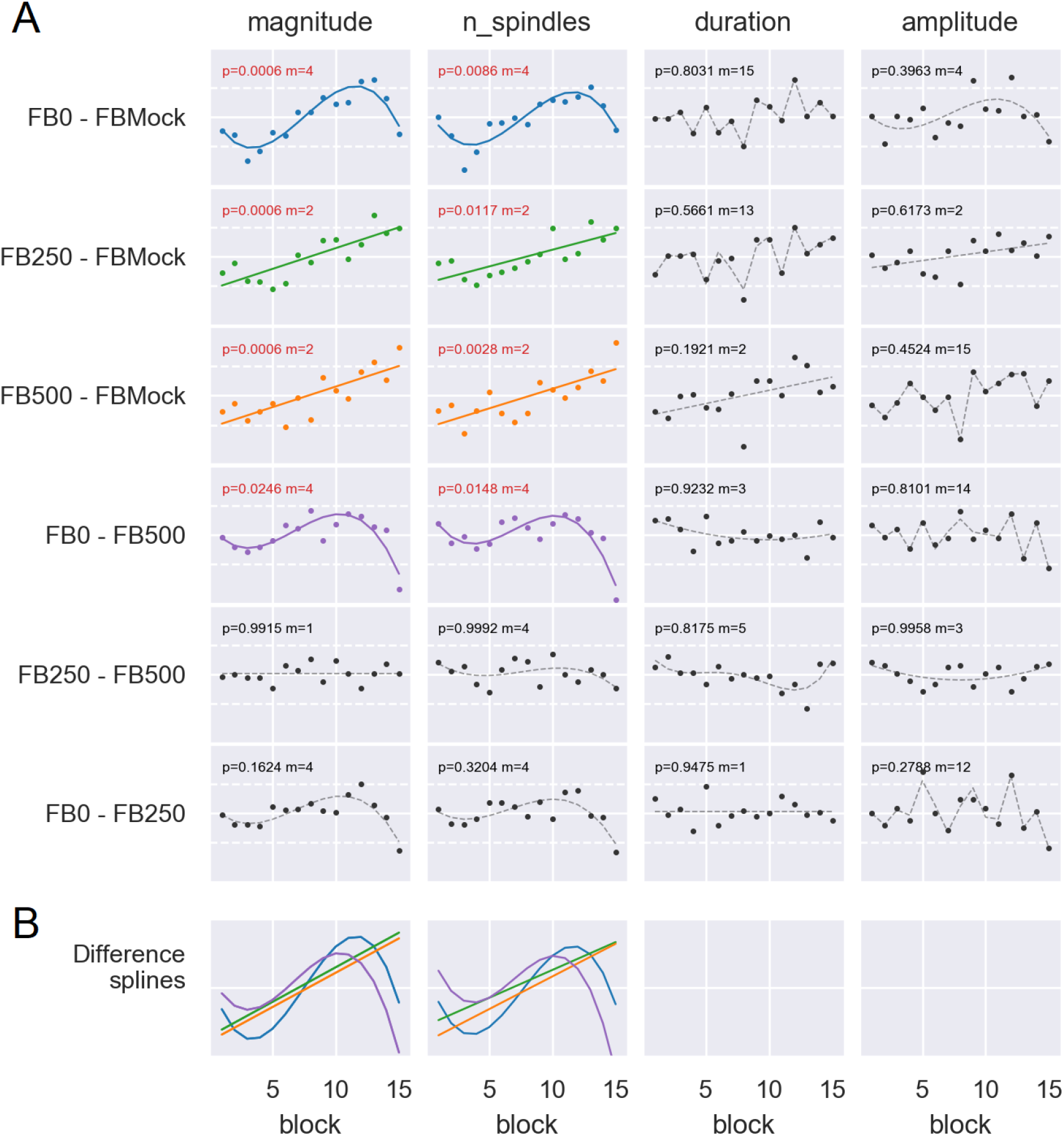
A) Pairwise *t*-statistics and AN-test results comparing dynamics of changes in the three morphological characteristics of alpha activity between all pairs of conditions for threshold factor *μ* = 2.5. *p*-values are FDR corrected. To facilitate comparison we have also included magnitude dynamics as the left-most column B) Superimposed parametric representations of the statistically significant difference between the compared learning curves. The difference exists only in magnitude and the incidence rate parameters.

Reassuringly, the S-shaped profiles observed in the results for burst magnitude (the first column) for FB0-FBMock and FB0-FB500 pairs replicate the profiles present in the incidence rate curves (the second column) and the linear trends observed for the other pairs are also very similar when magnitude and incidence rate data are compared. Finally, both magnitude and the incidence rate curves differ when FB0 and FB500 conditions are compared, and an S-shaped differential profile is revealed (*p* = 0.0237, FDR corrected). As evident from Figure 7 B that shows superimposed parametric profiles for the statistically significant difference, FB0 vs. FB500 and FB0 vs. FBMock shape difference contains a steep slope and a prominent decline at the end of the training contributed by FB0 condition.

The threshold value used for detection of alpha spindles could significantly affect the results described above. To explore the robustness of our findings, we repeated the above analyses for a range of threshold factor values *μ* ∈ [1, 3]. These results are summarized in Figure 8 where the FDR-corrected *p*-values obtained with AN-test comparing the corresponding pairs of conditions are color-coded. One can observe statistically significant variations in the learning curves for the incidence rate parameter for the majority of the explored thresholds when comparing true NFB conditions with mock NFB. Also, for a limited range of intermediate threshold values, significant changes are present in the shapes of the incidence rate learning curves for comparison of FB0 and FB500 conditions.

**Figure 8:**
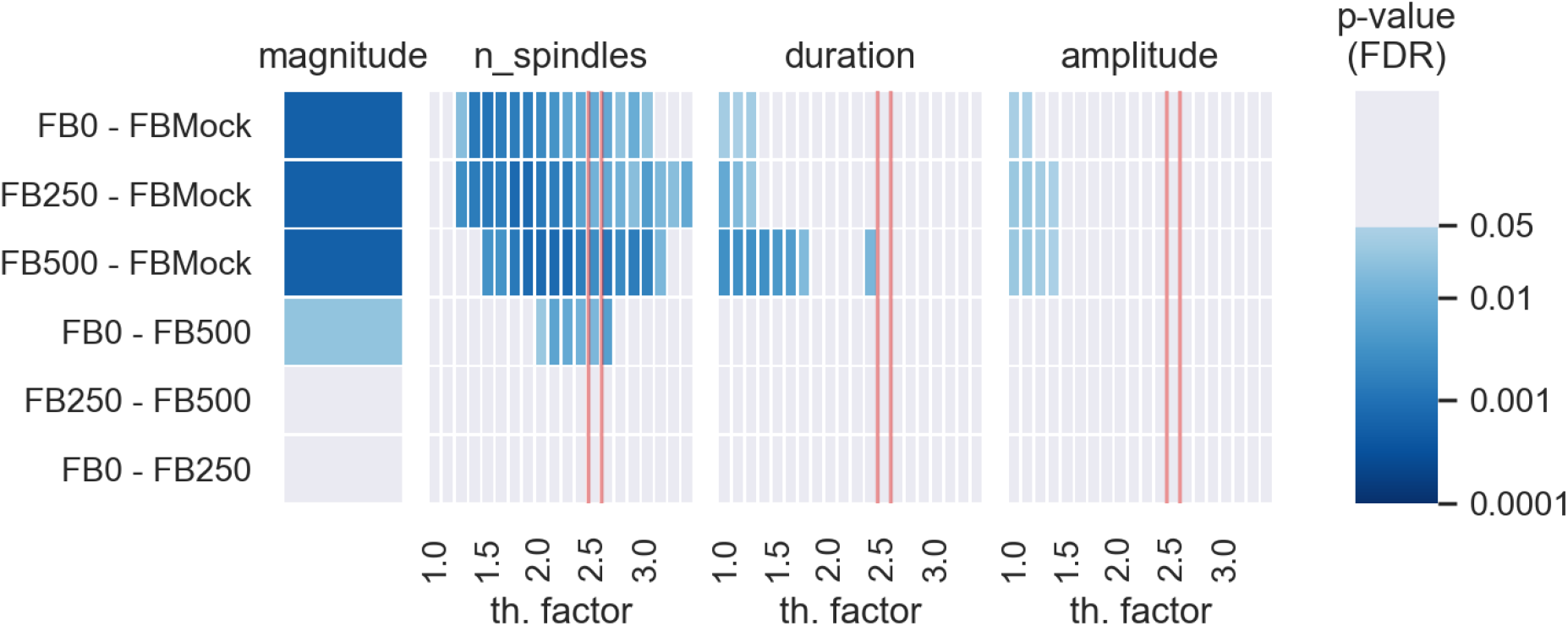
FDR corrected *p*-values for the pairwise AN-test exploring the effect of feedback latency on different morphological characteristics of the P4 alpha rhythm. Separate tests are performed for each threshold factor *μ* in a range from 1 to 3. The color indicates statistical significance based on *α* = 0.05 significance level. Red lines indicates results for optimal threshold factor *μ* = 2.5 corresponding to the minimal mutual information between the three morphological characteristics of the alpha-band activity

For a small range of thresholds that corresponds to low values of *μ* we consistently observe significant changes in burst duration and amplitude for the three true feedback conditions with respect to FBMock. These differences are stronger for FB500 condition where they are present for a broader range of threshold values. Interestingly, one can see a somewhat complementary pictures in comparisons between the three feedback conditions and mock feedback. For the subset of low threshold values we have observed differences in training dynamics for the amplitude and duration of alpha bursts whereas no difference is present in the incidence rate of alpha spindles. Yet, for the other(by far larger) contiguous subset of threshold values a difference is present in the incidence rate but not in burst amplitude and duration parameters.

### Increased latency negatively affects the magnitude of sustained changes induced by NFB

As illustrated in Figure 1 B, we recorded 2 minutes of resting-state baseline EEG activity before and after NFB training. Figure 9 A shows the changes in the mean alpha power between the two baseline intervals for all subjects included in the four groups. Only FB0 condition shows statistical tendency for the growth in post-intervention alpha magnitude, *p* = 0.022(0.0881), FDR corrected *p*-value is shown in brackets. The confidence interval for the mean paired difference in FB0 condition lies strictly in the positive range while the CIs for the other conditions include zero. No significant differences in feedback induced mean power gain was found via direct pairwise comparison of the three feedback conditions (FB0, FB250, FB500).

**Figure 9:**
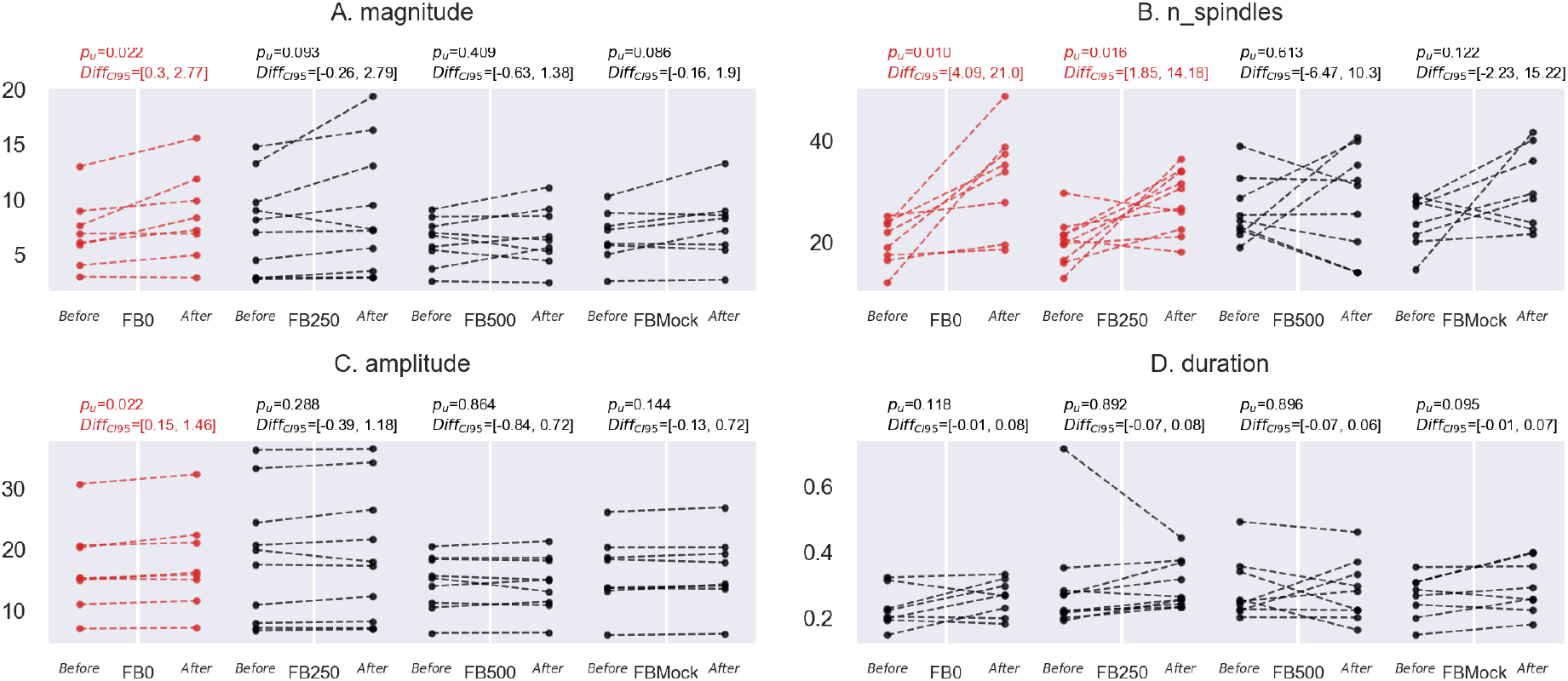
The pre-training and post-training baselines for four conditions of the study. A: Alpha magnitude. B: Burst incidence rate. C: Burst amplitude. D: Burst duration. The text above each graph indicates the 95% confidence interval for the mean paired difference and uncorrected *p*-values. FB0 condition has a significant increase in post-training alpha magnitude, incidence rate and a small but consistent growth in the burst amplitude parameter.

Next, we analyzed the sustained changes in the parameters of alpha spindles caused by NFB training. The results are summarized in panels B-D of Figure 9. In panel B, one can see an increase in spindles incidence rate in the post-training baseline as compared to the pre-training data: this effect is significant for FB0 and FB250 conditions. By visual inspection, the effect is stronger for FB0 and the CI of the difference is shifted to the right as compared to FB250 condition. One can also see a small but consistent and significant (*p* < 0.01) increase of burst amplitude for FB0 but not the other conditions. Therefore, we can conclude that the incidence rate of alpha spindles is a good indicator of learning during the ongoing training, as well of the lasting effects of NFB intervention. Interestingly and importantly, latency affects the extent to which NFB-induced changes persist after the training.

Most of what we have described above refers to within-group analysis or the analysis with respect to the mock neurofeedback condition. In order to demonstrate the effect of latency on the efficacy of NFB intervention more directly, we took advantage of the fact that feedback latency can be considered as a continuous variable. Accordingly, we performed a regression analysis, where statistical significance was quantified with parametric and non-parametric randomization tests. The results are presented in Figure 10 where one can see that only changes in the incidence rate significantly depended on NFB latency. The sustained changes were stronger for shorter latency.

**Figure 10:**
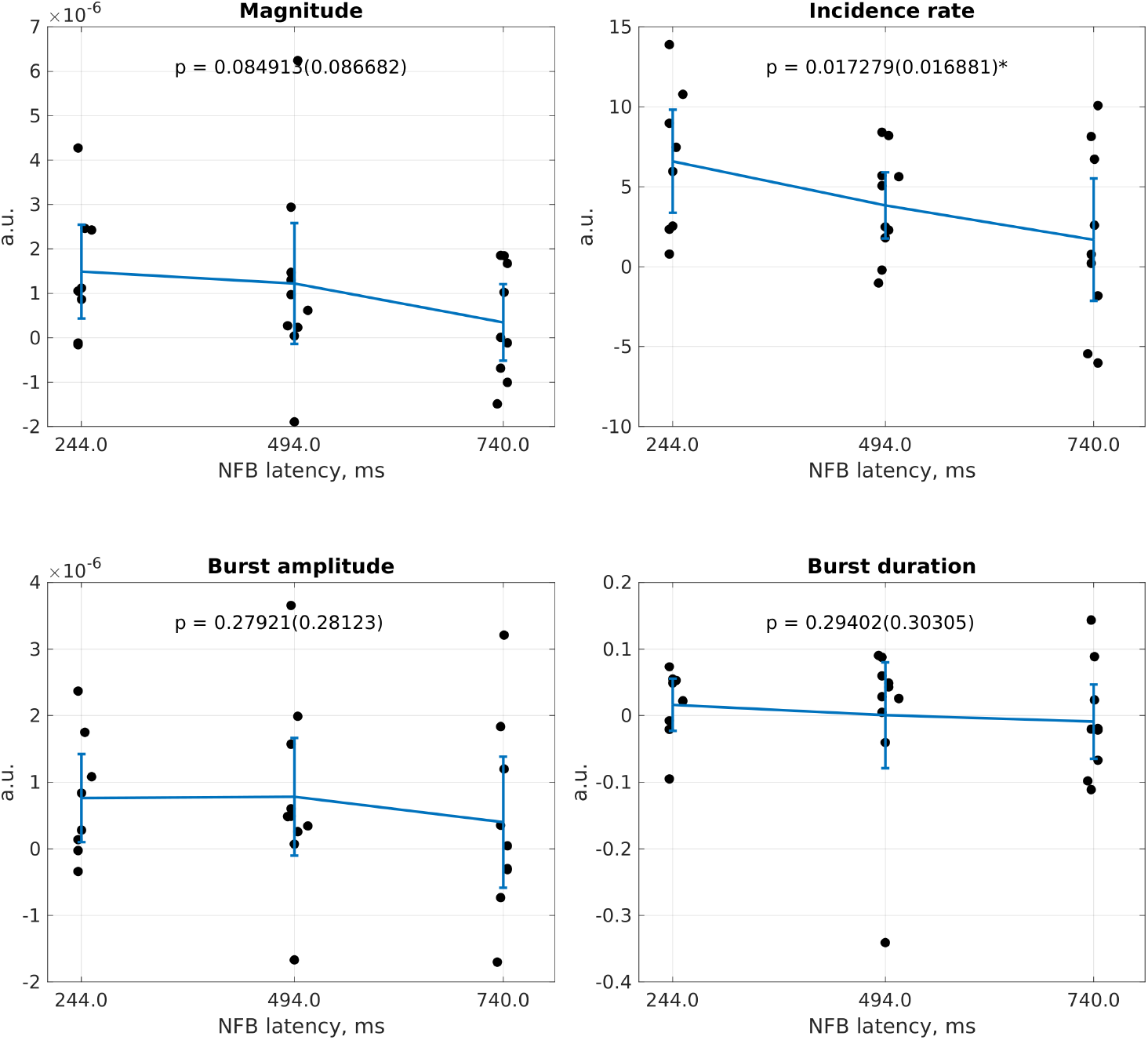
The magnitude of sustained changes in alpha rhythm parameters as a function of actual mean feedback latency for A) alpha magnitude, B)burst incidence rate, C) burst amplitude and D) burst duration. The text on each panel shows *p*-value for the hypothesis that the correlation coefficient is zero obtained with the parametric *t*-test and with a randomization test (in brackets). Only gain in alpha-spindles appear to have a significant linear dependence on feedback latency.

Thus, incidence rate of alpha spindles is the only parameter susceptible to NFB training, and it is the parameter that is affected when latency is varied, with clear improvement in training effects upon spindle incidence rate when latency is shortened. This result is consistent with the findings of our previous study where we investigated the effect of NFB training on spindle incidence rate, but did not vary latency [41]. Most important for practical applications, sustained changes in spindle incidence rate can be evoked only with short-latency NFB. We conclude that clinical alpha rhythm-based approaches should focus on shortening NFB latency and monitoring spindle incidence rate.

## Discussion

In this study, subjects were aided with NFB so that they could increase their parietal alpha activity. We implemented an experimental paradigm where NFB latency was precisely controlled and manipulated programmatically. The minimal latency that we could achieve with our hardware and software setup was the end-to-end delay of 244 ms, of which 100 ms corresponded to the causal estimation of the narrow-band EEG envelope [56]. We either kept this latency fixed or artificially imposed an extra delay. Additionally, a mock NFB condition was tested. The four experimental conditions (0, 250 and 500-ms added delays, and mock NFB) were run in separate groups of subjects.

During experimental planning, we reasoned that 10 subjects needed to be tested per group, making a total of 40 subjects. This is an appropriate sample for an exploratory/proof-of concept study, which is also consistent with the literature. Indeed, 10 subjects per group with test power of 0.8 would allow us to detect correlation of *ρ* = 0.7 between a parameter of interest and the training block number, which corresponds to correlation coefficient values observed in our previous study [41]. To explore more subtle effects, we took measures to increase the statistical power of our testing procedure. We employed a SNR based stratification procedure and used data normalization step to reduce inter-subject variability. Furthermore, we used the adaptive Neyman test and compared the learning curves which exploits their potential smoothness. When comparing baseline activity levels we used a paired test. Data from 5 subjects had to be removed because of the problems with EEG recordings (see Methods), which slightly decreased our sample compared to the original plan.

Our detailed study of the latency effects extends the previous work where cortical alpha rhythm served as the source of NFB [36], [1], [12], [19], [21], [61], [41]. In agreement with these previous studies, our subjects were able to increase the average magnitude of their alpha activity during 30 minutes of NFB training, and for the shortest NFB latency this increase was sustained after the training. By contrast, only a small increase in alpha activity was observed with mock NFB.

Consistent with our previous study [41], we observed clear changes in the incidence rate of alpha-activity bursts: as participants trained with NFB, these neural events became more frequent. Other changes in the structure of alpha-band activity, such as amplitude and duration of alpha bursts, were significantly less pronounced.

Interestingly, this result replicates our earlier findings [41] that showed, under different experimental settings, that NFB affects the incidence rate of alpha-spindles rather than influencing their shape. Thus, alpha spindles could be considered as discrete events whose probability changes as the result of NFB training. Relevant observations have been made regarding the beta-band transient events whose incidence rate, but not duration or amplitude predict motor performance across a range of species[54]. Thus, mean power in a particular spectral band – the parameter that has been traditionally used as a target parameter for NFB – crucially depends on the rate of occurrence of the discrete harmonic events. Therefore, we envision a NFB paradigm where these discrete short-lived events are specifically targeted by operant conditioning which in turn requires timely feedback presentation.

The goal of NFB training can be defined as attaining sustained changes in certain neural patterns. Therefore, the finding is important that only the lowest-latency NFB resulted in a sustained effect in our experiments. For this condition, average alpha amplitude was elevated, as evident from the comparison of baseline EEG recorded prior to NFB training with the EEG recorded after the training was completed (Figure 9.A). Incidence rate of alpha spindles was the major parameter that accounted for this change in baseline activity. Curiously, spindle rate increased not only for the shortest latency but also weakly for the condition where 250 ms were added to the base latency (Figure 9.B). In addition to the major effect of spindle rate, we observed a small but consistent and statistically significant change in the amplitude of alpha-spindles that occurred only for the shortest latency (Figure 9.C).

Several insights regarding the role of NFB latency can be gained from Figure 4. For the lowest latency, the initial rise in alpha activity was the steepest compared to the other conditions. Following the rise, alpha activity stabilized earlier and at a higher level when the latency was the shortest. Curiously, for the lowest latency, alpha activity declined over the last three training blocks. We used the adaptive Neyman test (AN-test) [15] useful to quantify this complex behavior. This test revealed an S-shaped profile for training with the lowest NFB latency whereas the dynamics for the longer latency were best described as linear trends that clearly differed from the trend for mock NFB. These results can be appreciated in Figure 5 that shows the time course of these changes and the splines fitted to *t*-statistics profiles *x_k_*, *k* = 1, …, 15 (see Appendix: Comparing learning curves for details). Spindle incidence rate practically mirrored the time course of the average alpha amplitude. By contrast, the curves for spindle amplitude and duration were not significantly different from those observed for mock NFB. The contribution of spindle rate was quite robust as it persisted for a broad range of threshold values that defined spindle events.

Moreover, the regression analysis performed on the data from 27 subjects trained on NFB with three different latencies showed a significant negative correlation between latency and the sustained gain in spindle incidence rate. This result directly demonstrates that NFB efficacy improves when latency is shortened.

Having established that NFB latency had a strong effect on spindle rate and little effect on the other parameters of alpha activity, we need to find an explanation for this finding. We suggest that our results could be explained by a reinforcement-learning mechanism, where NFB reinforces a neural pattern that coincides with NFB arrival, that is Hebbian plasticity is involved that strengthens a neural circuit that generates a particular activity pattern. Accordingly, the shorter the latency, the higher is the likelihood that NFB would reinforce the neural pattern that triggered the NFB, whereas a reinforcement that arrives with a lag would have a weaker effect on original neural pattern and the original circuit (i.e. a temporal discounting effect). Only with minimal NFB latency, robust changes in parietal alpha activity could be achieved. In this shortest-delay condition, NFB was initiated by an alpha spindle and arrived when the spindle was still ongoing, which reinforced the spindle and made its occurrence in the future more probable. Since an alpha spindle is a cortical event that has a stable structure (amplitude and duration), such reinforcement mechanism affects primarily the rate of this event but not its shape or duration. With the longer latencies of 250-500 ms, NFB is much less temporally specific and consequently less effective. Indeed, with the longer latency, NFB became less specific to the desired state transition to the oscillatory state simply because of increased likelihood that the original spindle has already completed by the time NFB arrives. Notably, participants in our experiments were asked to avoid any conscious strategy for modulating alpha activity. Under these conditions, learning was automatic and dependent on the Hebbian mechanism described above.

Overall, our findings suggest that NFB latency is a crucial parameter that needs to be minimized to achieve desired changes in the fine characteristics of EEG activity. While we experimented with relatively long NFB delays in this study, future work should examine shorter delays, particularly those on the order of 50 ms and lower. Such a low latency would enhance the sense of agency [14] and harness the power of automatic learning [30] by directly and specifically interacting with brain-state transitions. To achieve this desired latency decrease, more efficient signal processing pipelines are needed that use optimized hardware-software communication protocols, as well as more sophisticated signal processing pipelines for the extraction of oscillation parameters from brain activity [56, 34, 52].

## Supporting information

CRED-nf checklist

## Acknowledgements

This work is supported by the Center for Bioelectric Interfaces NRU HSE, RF Government grant, ag. No.14.641.31.0003.

## Appendix: Comparing learning curves

For every pair of FB groups, we compared learning curves between the two conditions and tested the null hypothesis (H0) about similarity of the learning curves. For such hypothesis testing, we used the adaptive Neyman test (AN-test) described in [15]. In contrast to the simple 2-sample T-test, AN-test takes into account the temporal connectivity between curve points. Also, in contrast to linear regression models, AN-test is not restricted to the linear learning curve prior and allows curves and the mean difference between the curves to be nonlinear. The steps of this test are the following:

1. For each pair of conditions {*i*, *j*} and for each block number *k* = 1..15 compute T-statistic *x_k_* to estimate the difference between average learning curves:

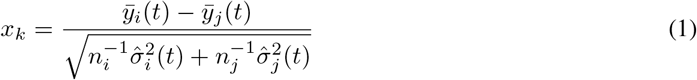
where 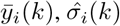 and 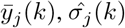, are the sample mean and sample standard deviation for the *k*-th training block in the *i*-th and *j*-th conditions, see also formula (13) in the original paper [15].
2. Represent vector **x** = (*x*_1_, …, *x*_15_) as a linear combination of predefined basis vectors **b**_*n*_ (*n*=1..15) that form transform matrix **B**. The corresponded coefficients of such decomposition can be obtained as 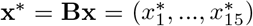. In the original paper this decomposition is obtained by Fourier transform. Here we used basis vectors **b**_*n*_ that corresponded to the discrete version of Legendre polynomials. This way the basis vectors describe the following temporal features: **b_1_** - *constant level*, **b_2_** - *linear trend*, **b_3_** - *U-shaped curve*, **b_4_** - *S-shaped curve*, etc. We expected that learning curves have more natural representation as the weighted sum of such temporal features rather than that formed with Fourier basis.
3. For the first *m* coefficients 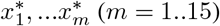 estimate final statistics (see formula (6) [15]). Find *m\** for which the statistic reaches maximal value 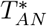. If 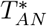 is too large than H0 can be rejected.

To determine the *p*-value we used finite sample distribution of 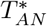 based on 200 thousand Monte-Carlo simulations. This distribution was obtained for each degree of freedom for the *T*-statistic in step 1. Note that the number of degrees of freedom is defined here as *df* = *n*_1_ + *n*_2_ – 2 where *n*_1_ and *n*_2_ is number of subjects in the first and second conditions in the pair. Additionally, we used value *m*\* from step 3 to recover smooth approximation 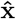 of learning curves difference by using the inverse transform 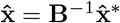, where 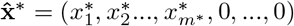 is a vector of length 15 which contains only first *m*\* non-zero coefficients.

