## Supplementary material for "Short-Delay Neurofeedback Facilitates Training of the Parietal Alpha Rhythm": CRED-nf checklist

### CRED-nf checklist summary

*27 July, 2020*

**Manuscript title:** Delayed neurofeedback impedes training of the parietal alpha-band power
**Corresponding Author:** Alexey Ossadtchi

| Item No. | Checklist item | Manuscript Details |
|---|---|---|
| **Pre-experiment** | | |
| 1a | Pre-register experimental protocol and planned analyses | *This experiment was not preregistered* |
| 1b | Justify sample size | During experimental planning, we reasoned that 10 subjects needed to be tested per group, making to a total of 40 subjects. This is an appropriate sample for an exploratory/proof-of concept study, which is also consistent with the literature. Indeed, 10 subjects per group with test power of 0.8 would allow us to detect correlation of r = 0.7 between a parameter of interest and the training block number, which corresponds to correlation coefficient values observed in our previous study (Ossadtchi et al. 2017). To explore more subtle effects, we took measures to increase the statistical power of our testing procedure. We employed a SNR based stratification procedure and used data normalization step to reduce inter-subject variability. Furthermore, we used the adaptive Neyman test and compared the learning curves which exploits their potential smoothness. When comparing baseline activity levels we used a paired test. Data from 5 subjects had to be removed because of the problems with EEG recordings (see Methods), which slightly decreased our sample compared to the original plan. We could not run additional experiments because of COVID-19 epidemic. |
| **Control groups** | | |
| 2a | Employ control group(s) or control condition(s) | The participants were split into four equal groups, each with different NFB settings: (1) NFB with no delay added to the system's base latency of 244 ms (FB0), (2) NFB with a 250-ms delay added (FB250), (3) NFB with a 500-ms delay added (FB500), (4) mock NFB where the feedback was generated from EEG data taken from a different participant (FBmock). |

| | | |
|---|---|---|
| 2b | When leveraging experimental designs where a double-blind is possible, use a double-blind | The experimental flow implied minimal interaction between the experimenter and the subject after the point when the group assignment was known as this happened using the described randomized stratification procedure past the moment when individual filters were setup. This means that briefing the subject, electrodes installation, and recording of the first two 1-minute baselines occurred before the group assignment was generated by the computer and saved in the subject's folder. |
| 2c | Blind those who rate the outcomes | The results of this stereotypical processing were grouped using the group assignment variable and the appropriate statistical comparisons were also performed automatically. |
| | Blind those who analyse the data | When processing the data and generating the results we analyzed all data with the scripts looping over all subjects from all four groups and applying the exactly identical processing pipeline. |
| 2d | Examine to what extent participants and experimenters remain blinded | *No measures were taken to examine whether participants and experimenters remained blind* |
| 2e | In clinical efficacy studies, employ a standard-of-care intervention group as a benchmark for improvement | *NA: This is not a clinical efficacy study* |
| **Control measures** | | |
| 3a | Collect data on psychosocial factors | The SNR was calculated before the main session from the one-minute baseline as the ratio of the average power spectral density (PSD) magnitude within the individually determined target frequency range to the mean PSD magnitude within the two flankers sub-bands. The participants with SNR less or equal to 1 were not included into the pool of subjects. The participants with SNR greater than 1 were assigned to one of the experimental groups with a stratified sampling procedure (see ”Composition of subject group” section above). |
| 3b | Report whether participants were provided with a strategy | All participants were given the same instruction. They were instructed to refrain from using any conscious strategy. This assured that learning to control the parietal alpha rhythm was automatic, which was the mode where the latency of a continuous NFB should hypothetically have a stronger effect on the performance. |
| 3c | Report the strategies participants used | Participants in our experiments avoided any conscious strategy for modulating alpha activity. |

| | | |
|---|---|---|
| 3d | Report methods used for online-data processing and artifact correction | The experiments started with the recording where a subject first looked at the fixation cross for one minute and then closed the eyes for one more minute. We then used these data to build a spatial filter based on the independent component analysis (ICA) to remove the artifacts caused by eye movements and blinking. This approach decomposed the EEG signals into independent components, including the ones containing the artifacts. Ocular artifact components were detected as those with the largest value of mutual information of their time-series and the signals in Fp1 and Fp2 channels, which are closest to the eyes. A spatial filter matrix was then constructed for the subsequent online application during the NFB sessions. Bandpass filters for extracting alpha activity were built separately for each individual. The signal was taken from channel P4. To detect alpha activity, we started with the frequency range 8 to 12 Hz, and then adjusted the range and filter parameters for each individual subject. We determined the central frequency of the rhythm by visual inspection and then set the range relative to the central frequency. With this method, the selected target signal frequency range accurately reflected the individual subject alpha rhythm properties. |
| 3e | Report condition and group effects for artifacts | *Condition and group effects for artifacts were not measured, or not reported in the manuscript* |
| **Feedback specifications** | | |
| 4a | Report how the online-feature extraction was defined | Alpha-rhythm envelope was extracted with the cFIR approach . In this method, the raw EEG signal is transformed into a narrowband analytic signal, a complex-valued function whose absolute value corresponds to an instantaneous amplitude(or envelope) of the rhythm. The cFIR method allows us to explicitly define NFB latency and obtain a more accurate envelope estimate for a specified latency compared to the other approaches frequently used for quantification of narrowband components in the EEG data. This speed-accuracy trade-off can be appreciated from the accuracy vs. processing delay curves for the cFIR and the commonly used approach based on narrow-band filtering followed by signal rectification. In the present study, we set the cFIR delay parameter to 100 ms . This setting corresponds to the correlation coefficient of 0.85 +- 0.1 between the off-line casually estimated and the on-line reconstructed envelopes. |
| 4b | Report and justify the reinforcement schedule | An alpha envelope was visualized as a circle with a pulsating outline. The subjects had to smooth that outline by increasing their P4 alpha-band power. |
| 4c | Report the feedback modality and content | An alpha envelope was visualized as a circle with a pulsating outline. |
| 4d | Collect and report all brain activity variable(s) and/or contrasts used for feedback, as displayed to experimental participants | The NFB signal was derived from the P4 channel (corresponding to the right parietal region). |

| | | |
|---|---|---|
| 4e | Report the hardware and software used | Subjects sat in a comfortable chair at a distance of 80 cm from an LCD monitor with a 24-cm diagonal and a 60-Hz refresh rate. These settings remained constant throughout the entire experiment. EEG signals were recorded using 32 AgCl electrodes placed according to the 10-20-system. Each EEG channel was sampled at 500 Hz using an NVX-136 amplifier (Medical Computer Systems Ltd) and bandpass-filtered in the 0,5 - 70 Hz band. These preprocessing filters incurred an overall delay of no more than 10 milliseconds for the bandwidth of interest (8–12 Hz). Digital common ear reference was derived from the electrodes placed on both ears. The impedance for each electrode was kept below 10 KOhm. |
| **Outcome measures - brain** | | |
| 5a | Report neurofeedback regulation success based on the feedback signal | Figure 4-9 |
| 5b | Plot within-session and between-session regulation blocks of feedback variable(s), as well as pre-to-post resting baselines or contrasts | Figure 4, Figure 6 |
| 5c | Statistically compare the experimental condition/group to the control condition(s)/group(s) (not only each group to baseline measures) | Figure 5, Figure 8, Figure 9 |
| **Outcome measures - behaviour** | | |
| 6a | Include measures of clinical or behavioural significance, defined a priori, and describe whether they were reached | *NA: the study does not take cognitive or behavioural measures* |
| 6b | Run correlational analyses between regulation success and behavioural outcomes | *NA: the study does not take cognitive or behavioural measures* |
| **Data storage** | | |
| 7a | Upload all materials, analysis scripts, code, and raw data used for analyses, as well as final values, to an open access data repository, when feasible | https://github.com/nikolaims/delayed_nfb |
